# Simultaneous and sequential multi-species coronavirus vaccination

**DOI:** 10.1101/2022.05.07.491038

**Authors:** Lei Peng, Zhenhao Fang, Paul A. Renauer, Andrew McNamara, Jonathan J. Park, Qianqian Lin, Xiaoyu Zhou, Matthew B. Dong, Biqing Zhu, Hongyu Zhao, Craig B. Wilen, Sidi Chen

**Affiliations:** Department of Genetics, Yale University School of Medicine, New Haven, CT, USA; System Biology Institute, Yale University, West Haven, CT, USA; Center for Cancer Systems Biology, Yale University, West Haven, CT, USA; Molecular Cell Biology, Genetics, and Development Program, Yale University, New Haven, CT, USA; Department of Immunobiology, Yale University, New Haven, CT, USA; Department of Laboratory Medicine, Yale University, New Haven, CT, USA; M.D.-Ph.D. Program, Yale University, West Haven, CT, USA; Immunobiology Program, Yale University, New Haven, CT, USA; Computational Biology and Bioinformatics Program, Yale University, New Haven, CT, USA; Department of Biostatistics, Yale University School of Public Health, New Haven, CT, USA; Yale Comprehensive Cancer Center, Yale University School of Medicine, New Haven, CT, USA; Yale Stem Cell Center, Yale University School of Medicine, New Haven, CT, USA; Yale Center for Biomedical Data Science, Yale University School of Medicine, New Haven, CT, USA

**Keywords:** Lipid nanoparticle mRNA vaccine, vaccination schedule, multiplexed vaccination, sequential vaccination, simultaneous vaccination, multi-species coronavirus vaccine, COVID-19, SARS-CoV-2, SARS-CoV, MERS-CoV, cross-reactivity, systems immunology, single cell profiling

## Abstract

Although successful COVID-19 vaccines have been developed, multiple pathogenic coronavirus species exist, urging for development of multi-species coronavirus vaccines. Here we developed prototype LNP-mRNA vaccine candidates against SARS-CoV-2 (Delta variant), SARS-CoV and MERS-CoV, and test how multiplexing of these LNP-mRNAs can induce effective immune responses in animal models. A triplex scheme of LNP-mRNA vaccination induced antigen-specific antibody responses against SARS-CoV-2, SARS-CoV and MERS-CoV, with a relatively weaker MERS-CoV response in this setting. Single cell RNA-seq profiled the global systemic immune repertoires and the respective transcriptome signatures of multiplexed vaccinated animals, which revealed a systemic increase in activated B cells, as well as differential gene expression signatures across major adaptive immune cells. Sequential vaccination showed potent antibody responses against all three species, significantly stronger than simultaneous vaccination in mixture. These data demonstrated the feasibility, antibody responses and single cell immune profiles of multi-species coronavirus vaccination. The direct comparison between simultaneous and sequential vaccination offers insights on optimization of vaccination schedules to provide broad and potent antibody immunity against three major pathogenic coronavirus species.

**One sentence summary:** Multiplexed mRNA vaccination in simultaneous and sequential modes provide broad and potent immunity against pathogenic coronavirus species.

## INTRODUCTION

*Coronaviridae* is a large family of viral species constantly evolving ^1^. Coronaviruses are genetically diverse RNA viruses that exhibit broad host range amongst mammals, where the infections cause a wide range of diseases, ranging from common cold to severe illnesses and death ^1 2^. Multiple zoonotic coronavirus species evolved to infect humans, became highly contagious, pathogenic and even fatal, causing worldwide pandemic ^1^. To date, seven known coronavirus species evolved to infect humans ^1^. There are three known highly pathogenic human coronavirus species to date, severe acute respiratory syndrome coronavirus (SARS-CoV), middle east respiratory syndrome coronavirus (MERS-CoV) and severe acute respiratory syndrome coronavirus 2 (SARS-CoV-2), all of which can cause severe respiratory or multi-organ diseases and can be fatal ^3^. There are also thousands of potentially highly pathogenic coronaviruses circulating in animal reservoirs globally ^2, 4–6^. SARS-CoV-2 is the pathogen that cause coronavirus disease 2019 (COVID-19) ^1^, an ongoing multi-wave worldwide pandemic ^7^ that claimed over 5 million lives to date. SARS-CoV and MERS-CoV emerged in humans in 2002 ^8^ and 2012 ^9^, and have high case fatality rates (∼10% for SARS-CoV and ∼35% for MERS-CoV, relative to ∼1% for SARS-CoV-2) ^10^. Thus, it is important to develop effective vaccines against these highly pathogenic coronavirus species. Before the COVID-19 pandemic, no effective vaccine had been approved to prevent spread of coronaviruses. Previous SARS and MERS vaccine devolvement ^11–15^, although at earlier stages, together with global efforts, led to rapid development of multiple COVID-19 vaccines against SARS-CoV-2 ^16^. The most prominent and efficacious vaccine belong to the lipid nanoparticle (LNP) mRNA vaccine category, with the first two emergency use approval issued to Moderna and Pfizer-BioNTech mRNA vaccines ^17, 18^. Although successful vaccines against SARS-CoV-2 have been developed to control COVID-19, no effective vaccines exist that can counter multiple pathogenic coronavirus species including SARS-CoV and MERS-CoV. Thus, it is important to develop new multi-species coronavirus vaccines, not only to help fight the ongoing pandemic, but also to prevent reemergence of these previously existed dangerous pathogens, as well as to gain insights to prepare for future zoonotic pathogenic coronavirus outbreaks.

The success of LNP-mRNA vaccine against COVID-19 led to the natural hypothesis of multiplexed vaccination against multiple coronavirus species. However, to date, there has been no reported approved or clinical stage vaccine specifically generated to target two or more of these highly pathogenic species. Even with such vaccine candidates generated, a number of important questions need to be answered. Certain prior studies in other virus families such as Influenza, HSV and CMV demonstrated initial feasibility of using two or more mRNA vaccine constructs in mixture ^19–21^. However, it is important to test multiplexing of several mRNAs directly for coronavirus species, which has yet to be done to date. Moreover, the immunogenicity of multi-species coronavirus vaccines need to be studied, for example, LNP-mRNA vaccines against MERS-CoV, SARS-CoV-2 and SARS-CoV. A recent study used a chimeric mRNA as vaccine candidates against two or more coronaviruses ^22^. However, chimeric spikes would not be able to capture all full-length spikes, losing part of critical antigenic regions (e.g. S1 or S2) for one species or the other. The two highly successful approved mRNA vaccines used full-length spike ^17, 18^. Finally, the optimal vaccination schedule needs to be explored involving a multiplexed vaccination, for example, would vaccination by administering all mRNAs simultaneous be effective, and if spacing out the different mRNA vaccine shots perform better. In order to gain answers to these important questions, we directly generated species-specific LNP-mRNA vaccine candidates and tested them in combination and in sequence *in vivo*. We generated LNP-mRNAs specifically encoding the HexaPro engineered full-length spikes of SARS-CoV-2 Delta variant, SARS-CoV and MERS-CoV, and systematically studied their immune responses in animal models.

## RESULTS

### Design and biophysical characterization of triplex coronavirus vaccine against SARS-CoV-2, SARS-CoV and MERS-CoV

We first designed vaccine candidate constructs encoding full-length spike mRNA of SARS-CoV-2 (labeled as **SARS2** for short) Delta variant (**Delta**), SARS-CoV (**SARS**) and MERS-CoV (**MERS**) (**Figure 1A-B; Figure S1**). Each construct contains a 5’untranslated region (UTR), an open reading frame (ORF), a 3’UTR and a polyA signal. The ORFs encode full-length spikes of defined species (SARS2, SARS and MERS), in which 6 additional proline mutations (HexaPro) were introduced in the S2 domain of the respective species (Figure 1a-b), based on the homologous amino acid positions of SARS-CoV-2, in order to improve expression and stable prefusion state of spikes ^23^. The Delta construct ORF encodes the spike of SARS-CoV-2 Delta variant, which has nine mutations (T19R, 156del, 157del, R158G, L452R, T478K, D614G, P681R, and Q1071H) as compared to the original “wildtype” virus (**WT**, **WA-1 or WA1**) virus (**Figure 1A-B**). To multiplex these constructs, we prepared equal-mass mixture of spike mRNA of Delta, SARS and MERS, which were then encapsulated by lipid nanoparticles on a microfluidics instrument, to generate a triplex LNP-mRNA formulation of vaccine candidate (termed as Triplex-CoV, or Triplex) (**Figure 1C**). We also generated a Delta singlet LNP-mRNA for testing in parallel. The size and homogeneity of assembled LNPs were evaluated by dynamic light scatter and transmission electron microscope (**Figure 1D-E**). The Delta LNP-mRNA and Triplex LNP-mRNA showed monodispersed size distribution with averaged radius of 70 ± 3.8 nm and 71 ± 3.6 nm, and polydispersity indices of 0.160 and 0.157, respectively. We tested each of these mRNA constructs and showed that they all successfully generate functional protein upon introduction into mammalian cells, as evident by surface binding to the cognate human receptors, hACE2 for SARS-CoV and SARS-CoV-2, and hDPP4 for MERS, respectively (**Figure 1F**).

**Figure 1.**
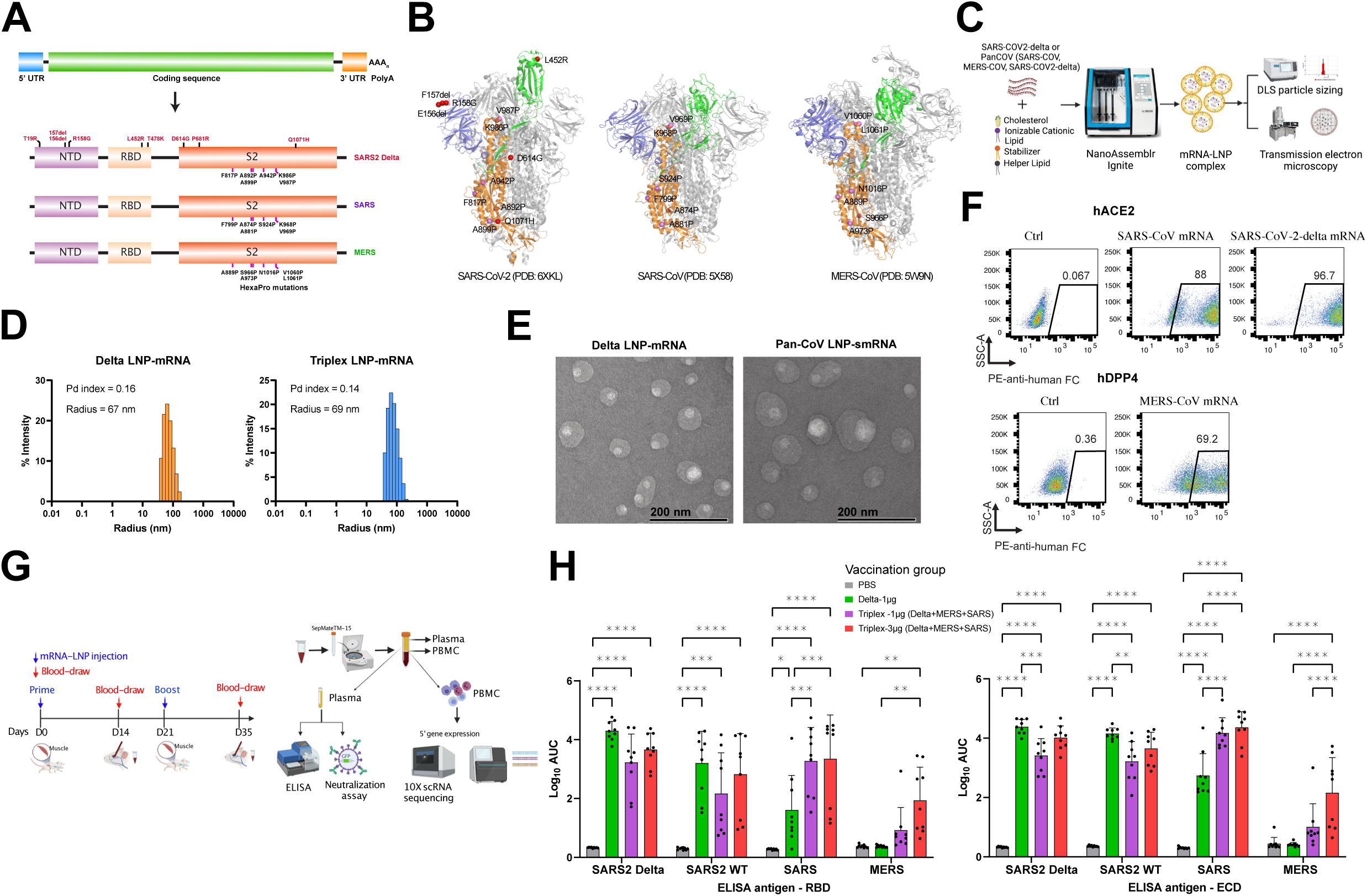
Antibody responses and single cell immune profiles of triplex and sequential LNP-mRNA vaccination against SARS-CoV-2 Delta, SARS-CoV and MERS-CoV *in vivo*. **(A)** Schematics of mRNA vaccine construct design against pathogenic human coronavirus species. Each construct has regulatory elements (5’UTR, 3’UTR and polyA) and spike ORF. The domain structures as well as engineered mutations of translated spike proteins of SARS-CoV-2 Delta variant (Delta), SARS-CoV (SARS) and MERS-CoV (MERS). **(B)** Engineered mutations in spike protein structures of SARS-CoV-2 Delta, SARS-CoV and MERS-CoV. The N-terminal domain (NTD, blue), receptor binding domain (RBD, green) and S2 subunit (orange) of one protomer along with homologous HexaPro mutations (pink) and Delta variant mutations (red) were highlighted in the spike trimer structures. **(C)** Schematics of characterization of LNP-mRNA vaccine formulations. Assembly procedure of LNP-mRNA vaccine on NanoAssemblr Ignite and downstream biophysical characterization assays. **(D)** Histogram displaying radius distribution of LNP-mRNA formulations of SARS-CoV-2 Delta and a Triplex (Delta + SARS + MERS) (abbreviated as Triplex-CoV or Triplex), measured by dynamic light scattering (DLS). The polydispersity index and mean radius of each LNP sample were shown at top left corner. **(E)** Transmission electron microscope (TEM) images of Delta and Triplex-CoV LNP-mRNAs. **(F)** Surface expression of functional spike proteins in 293T cells after electroporation of corresponding mRNA, as detected by human ACE2 or human DPP4 Fc fusion protein bound to PE anti-Fc antibody. **(G)** Schematics of vaccination schedule of the Triplex LNP-mRNA formulations, as well as downstream assays to evaluate the antibody responses and other immunological profiles. **(H)** Binding antibody titers of plasma samples from mice administered with PBS or different LNP-mRNAs (n = 9) against RBD or ectodomain (ECD) of SARS-CoV-2 wild type (WT, Wuhan/WA-1), Delta variant, SARS and MERS spikes. The binding antibody titers were quantified by area under curve of log_10_-transformed titration curve (log_10_ AUC) in Figure S2. The mice were intramuscularly injected with two doses (x2, 2 weeks apart) of PBS, 1μg SARS-CoV-2 Delta variant LNP-mRNA (delta), 1μg or 3μg equal mass mixture of Delta, SARS and MERS LNP-mRNA (Triplex-CoV). **Notes:** In the dot-box plots of this figure, each dot represents data from one mouse. Data are shown as mean ± s.e.m. plus individual data points in plots. Two-way ANOVA with Tukey’s multiple comparisons test was used to assess statistical significance. Statistical significance labels: * p < 0.05; ** p < 0.01; *** p < 0.001; **** p < 0.0001. Non-significant comparisons are not shown, unless otherwise noted as n.s., not significant.

### Immune responses to triplex coronavirus LNP-mRNA vaccination against SARS2, SARS and MERS

To evaluate the immunogenicity of Delta and Triplex LNP-mRNA vaccines, C57BL/6Ncr (B6) mice were immunized intramuscularly with two doses (prime and boost) of 1µg Delta LNP-mRNA, 1 µg or 3 µg (total mRNA mass) Triplex LNP-mRNA, three weeks apart (**Figure 1G**). The peripheral blood mononuclear cells (PBMCs) and plasma were collected two weeks post boost. The mice humoral response including binding and antibody response against spike antigens were examined using collected plasma samples. Single cell RNA-sequencing (scRNA-seq) was performed to profile the systemic immune repertoires and their respective transcriptomics in vaccinated animals (**Figure 1G**). Compared to the PBS control group, the 1µg Delta LNP-mRNA, 1µg and 3µg Triplex LNP-mRNA all elicited potent antibody response, as seen in the high post-boost binding antibody titers against both RBD and ECD of Delta, WT and SARS spikes (**Figure 1H; Figure S2; Figure S3A**). Among the three vaccination groups, only 3µg Triplex LNP-mRNA significantly boosted mice immunity to MERS antigens (**Figure 1H**). As the Delta and Triplex vaccines used the Delta variant as spike antigen, their responses to Delta ELISA antigen were found slightly higher than WT antigen (**Figure 1H**). Despite of the lack of SARS spike antigen in the vaccine, the Delta LNP-mRNA induced antibodies that cross-react with SARS spike, but not MERS spike (**Figure 1H**), consistent with the respective degree of homology between these species (**Figure S1**). The titers are at similarly high level between the 1µg and 3µg Triplex groups for SARS and SARS2 spikes (**Figure 1H**), while there is a trend of dose-dependent increase although statistically insignificant (**Figure 1H; Figure S2**). Compared to those of Triplex 1µg or 3µg groups, SARS-binding antibody titer in 1µg Delta LNP-mRNA group was significantly lower. A dose-dependent increase trend of antibody titers was observed for MERS spike in the two Triplex vaccination groups (**Figure 1H**). Considering Delta spike mRNA at the same dose, mice in the 1µg Delta and 3µg Triplex (that also contains 1µg Delta mRNA) groups showed similar titers of antibodies against SARS2 WT and Delta spikes, although an insignificant trend of lower titers was observed in the 3µg Triplex mice (**Figure 1H; Figure S2**). Both ECD and RBD ELISA antigen panels showed highly correlated results among four spike types used (**Figure S3B**). In addition, a subset of animals showed relatively higher titer to ECD than RBD (**Figure S3B,** off-the-diagonal data points), potentially due to the additional antibody reactivity outside RBD in those animals.

### Single cell immune repertoire mapping of multiplexed LNP-mRNA vaccinated animals

In order to gain insights on the global composition and transcriptional landscape of the immune cells, we performed single cell RNA-seq (scRNA-seq, scGEX) for immune-transcriptomics on the PBMC samples of Delta and Triplex LNP-mRNA vaccinated animals. As visualized in an overall Uniform Manifold Approximation and Projection (UMAP), from a total of 12 animals from 4 vaccination groups (Delta 1 µg, Triplex-CoV 1 µg and 3 µg dose groups), plus a placebo control group (PBS), we sequenced the transcriptomes of a total of 91,526 single cells (**Dataset S1**), which were visualized in reduced dimensional space by UMAP and clustered to identify cell population structure (**Figure 2A-B; Figure S3**). sing the expression of a set of canonical cell type specific markers, we identified 21 cell clusters as distinct immune cell populations (**Figure 2A-D; Figure 3A; Figure S3**). In this dataset, the identified cell clusters include various subsets of B lymphocytes (naïve B cell, activated B cell, unswitched memory B cell, switched memory B cell, pre-plasmablast, plasmablast and plasma cell); T lymphocytes of various subsets (naïve CD8 T cell, CD8 T effector, CD8 central memory T cell (Tcm), CD8 effector memory T cell (Tem), naïve CD4 T cell, Th1 type CD4 T cell, Th2 type CD4 T cell, regulatory T cell (Treg)); dendritic cells (DCs) of various subsets (pDC, cDC1, cDC2); as well as other immune cells (natural killer (NK) cell, macrophage and monocytes) (**Figure 2A; Figure 3B**). These immune cell populations have distinct gene expression signatures that clearly defined each population against others (**Figure 2B-C**), for example, distinct expression (in terms of both mean expression level and percentage in cluster) of *Cd19+H2-Aa+Ighd+Fcer2a+Cd27-*defines activated B cells; *Cd9+Sdc1+Cd19-Pax5-lo* defines plasma cells (**Figure 2A; Figure 3A-B; Figure S3**). Similarly, for T cell subset examples, *Cd3d+Cd4+Tbx21+Gzmb+* marks Th1 CD4 T cells; *Cd3d+Cd4+Foxp3+Il2ra+* marks Tregs; *Cd3d+Cd8b1+Ccr7+Cd44-Tcf7+* defines naïve CD8 T cells; *Cd3d+Cd8b1+Tcf7-Cd44+Ccr7-*defines CD8 effector T cells; *Itgam+Itgax+Cd24a-Sirpa+* defines cDC2 cells; *Itgam*-*Itgax*+*Bst2*+*Siglech*+ defines pDC; *Ncr1* defines NK cells; and *Itgam*+*Csf1r+Cd14+* defines monocytes (**Figure 2A, C, D; Figure 3A-C**). We then quantified the fractions of each cell type in each sample, to reveal a full picture of immune cell compositions in all vaccination groups profiled (**Figure 2E**). With these quantitative fractions, we then compared the systemic immune cell compositions between placebo and vaccinated animals (**Figure 2E**). While most of the clusters did not show significant difference in a gross cell population level, three populations (activated B cells, unswitched memory B cells and NK cells) showed significant differences between groups (**Figure 2E**). Interestingly, Triplex LNP-mRNA at both high and low doses of vaccination showed significantly increased level of activated B cell populations compared to both PBS and to Delta groups (**Figure 2E**). Both activated and memory B cell populations have been previously implicated for their important roles in SARS-CoV-2 immunity ^24–26^.

**Figure 2.**
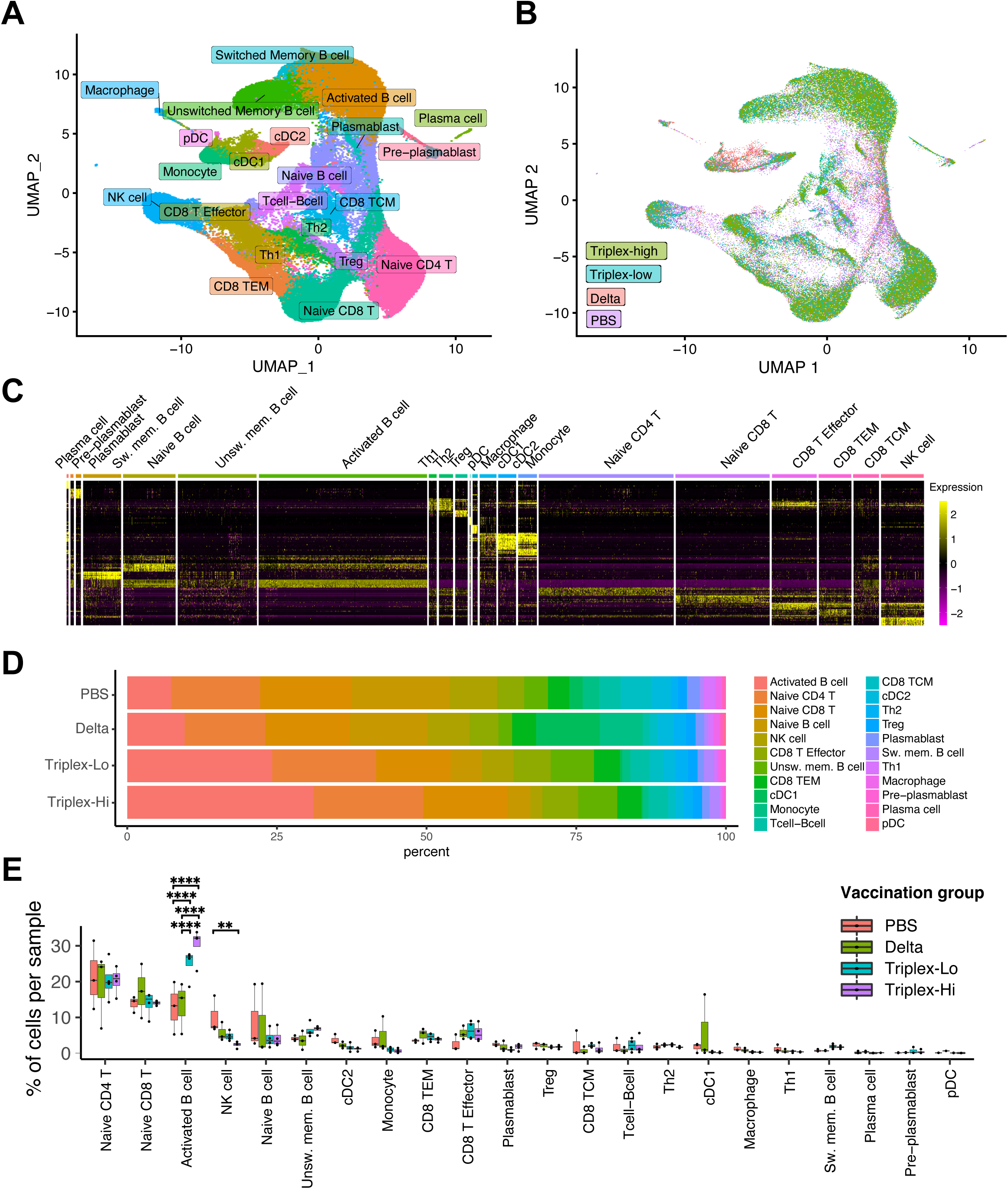
Single cell transcriptomics of animals vaccinated by multiplexed LNP-mRNA vaccine against SARS-CoV-2, SARS-CoV and MERS-CoV in mice. **(A)** UMAP visualization of all 91,526 cells pooled across samples and conditions. All identified clusters are shown with cell identities assigned, based on the expression of cell type specific markers. **(B)** Dot-whisker plot of immune cell proportions by cell type for each vaccination group: PBS, Delta, Triplex-lo and Triplex-hi; n = 3 mice each group. **(C)** Schematics of sequential vs. mixture vaccination schedules and sampling. In the **Sequential** vaccination schedule, vaccinations of SARS-CoV-2 Delta, MERS-CoV, and SARS-CoV were given in sequence separated by 3 weeks, each with 1μg LNP-mRNA prime and 1μg LNP-mRNA boost 3 weeks apart. In the **Mixture** vaccination schedule, vaccinations of SARS-CoV-2 Delta, MERS-CoV, and SARS-CoV were given simultaneously, each at 1μg LNP-mRNA (3μg total) for both prime and boost. The first dose and the blood sample harvest were done at the same day for both sequential and mixture schedules for comparison. **(D)** Dot-box plots summarizing binding antibody titers of plasma from mice administered with PBS, Sequential or Mixture LNP-mRNA vaccinations (n = 4 each) against RBD or ECD of SARS-CoV-2 WT/WA-1 and Delta variant, as well as SARS and MERS spikes. **(E)** Neutralization of plasma from mice treated with PBS, Sequential or Mixture LNP-mRNA vaccinations (n = 4 each); all tested against WT/WA-1 and Delta SARS-CoV-2, SARS-CoV and MERS-CoV pseudoviruses. The percent of GFP positive cells reflected the infection rate of host cells by pseudovirus and was plotted against the dilution factors of mice plasma to quantify neutralizing antibody titers. Neutralizing antibody titers in the form of reciprocal IC50 derived from fitting the titration curves with a logistic regression model. Each dot represents data from one mouse (n = 4 mice per group).

**Figure 3.**
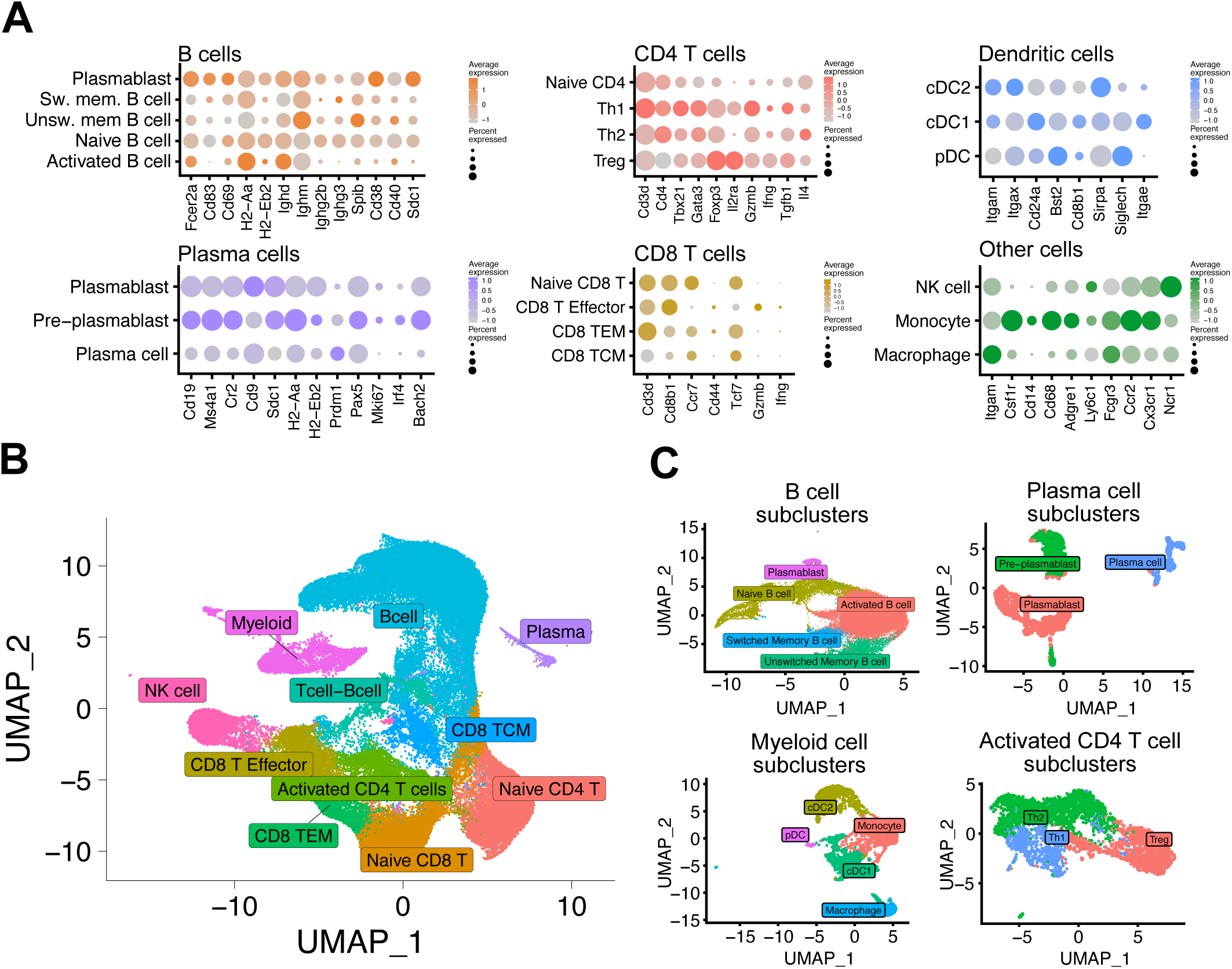
Single cell transcriptomics visualization, clustering and identification of major adaptive immune populations. **(A)** Bubble plots showing cell population clusters and their respective feature markers, showing expression of representative cell type-specific markers in T cells, NK cells, myeloid cells, B cells, and plasma cells. **(B)** UMAP clustering, color-coded by major immune cell populations including T cells, NK cells, myeloid cells, B cells, and plasma cells. **(C)** UMAP visualizations of sub-clustering of major immune cell populations for differential gene expression, including pooled B cells, plasma cells, myeloid cells, and activated CD4 T cells. **Notes:** Each indicated cell type in the analysis represents the pooled activated immune cell subsets from the overall UMAP: B cell = activated B cells, switched memory B cells, and unswitched memory B cells; CD4 T cells = Th1, Th2 and Treg; CD8 T cells = CD8 effector T cells, CD8 TEM, and CD8 TCM.

### Transcriptomic signatures of B and T cell populations of Triplex LNP-mRNA vaccinated animals

To examine the transcriptomic changes in the immune cell sub-populations upon vaccination, we performed differential expression (DE) analysis in the matched sub-populations between PBS and the several LNP-mRNA groups. We focused on the major adaptive immune cell populations, i.e. the pan activated B cell population (including all identified activated B cell subsets, merged as “B cell”), pan activated CD4 T cell population (all identified activated CD4 T cell subsets, “CD4 T cell”) and pan activated CD8 T cell population (all identified activated CD8 T cell subsets, “CD8 T cell”). Vaccination caused substantial transcriptome changes in the host animals’ B cells, CD4 T cells and CD8 T cells, as evidenced by the differential gene expression from vaccinated (Delta, Triplex-CoV/Triplex low and high dose groups) as compared to the PBS group (**Dataset S1; Figure 3C; Figure 4A; Figure S5; Figure S6**). To gain a broad, unbiased view of these transcriptomic changes, we performed a series of gene set and pathway analyses. These analyses revealed a number of altered pathways in the vaccinated animals B cells, CD4 T cells and CD8 T cells as compared to the PBS group (**Dataset S1; Figure 4A-B**). Network analysis of enriched pathways of differentially expressed genes highlighted the most significantly enriched member pathways (as meta-pathway), for the main adaptive immune cell types (B and T cells), for the three vaccination groups (**Figure 4C**). A top enriched pathway of the differentially expressed genes in B cells is B cell activation, where all three vaccines induced a higher expression of these genes (**Figure 4C**). In CD4 and CD8 T cells, common gene sets are observed, including immune system processes, immune cell differentiation, and T cell activation, consistent with the expected induction from vaccination (**Figure 4C**). Interestingly, in T cells, in the differentially expressed genes in all three vaccines, besides regulation of T cell activation and immune responses, basic metabolic pathways are also enriched, especially those involved in core metabolic functions such as oxidative phosphorylation and mitochondria respiratory chain activities (**Figure 4C**), consistent with the expectation that T cells are metabolically active upon vaccination. The Triplex vaccination induced strong B cell activation pathway clusters in B cells, as well as immune cell differentiation and metabolic activity gene sets in T cells (**Figure 4C**). These data reveal the broad gene expression signatures at the pathway and cluster levels, across the main adaptive immune cells (B and T cells) for the three vaccination groups. The transcriptomic signatures are largely coherent with the literature that these pathways are important for immunity against coronavirus infection and host defense ^27, 28^, as well as vaccine-induced immune responses ^29, 30^. These data revealed meta-pathway level gene expression changes in the B and T cells’ transcriptomes of the animals receiving multiplexed vaccination.

**Figure 4.**
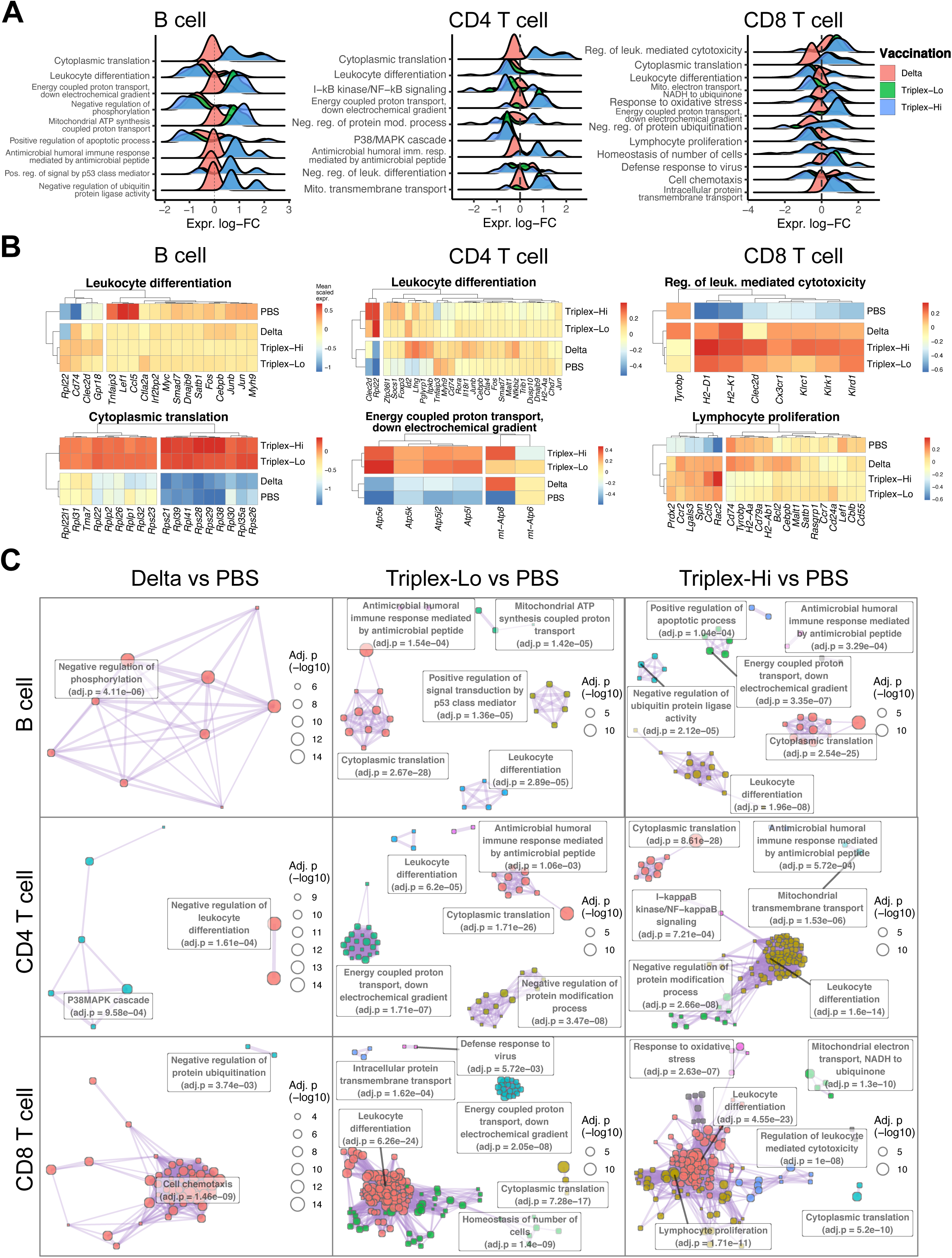
Differential expression, pathway signature and gene set cluster analyses of single cell transcriptomics for animals vaccinated by multiplexed LNP-mRNAs. **(A)** Ridge density plots showing the expression log fold change meta-pathway genes between different vaccination groups in different cell types. Each plot presents the top five meta-pathways in either Triplex-vs-PBS analysis, and only differentially expressed genes of either analysis were selected for each meta-pathway ridgeplot. Each indicated cell type in the analysis represents the pooled activated immune cell subsets from the overall UMAP: B cell = activated B cells, switched memory B cells, and unswitched memory B cells; CD4 T cells = Th1, Th2 and Treg; CD8 T cells = CD8 effector T cells, CD8 TEM, and CD8 TCM. **(B)** Heatmaps of differentially expressed genes between different vaccination groups of representative pathways in different cell types. **(C)** Network plots of enriched pathways of differentially expressed genes between the vaccination groups and PBS, in different cell types. **Notes:** Each dot is a pathway with the size and color representing the -log10 adjusted p value and the pathway cluster, respectively. Clusters are labeled with the most significantly enriched member pathway (meta-pathway). Colored representative meta-pathway clusters correspond to the colored text boxes.

### Direct comparison of sequential vs. simultaneous vaccination schedules for LNP-mRNA vaccination against three species

As observed above, Triplex LNP-mRNA vaccination is associated with reduction of antibody responses, we hypothesized that splitting such vaccination into separate doses may be a strategy to mitigate this loss of effectiveness. We therefore sought to perform a sequential vaccination schedule and test it in parallel with simultaneous vaccination with mRNAs in mixture (**Figure 5A**). In the **Sequential** vaccination schedule, vaccinations of SARS-CoV-2 Delta, MERS-CoV, and SARS-CoV were given in sequence separated by 3 weeks, each with 1μg LNP-mRNA prime and 1μg LNP-mRNA boost 3 weeks apart. In the **Mixture** vaccination schedule, vaccinations of SARS-CoV-2 Delta, MERS-CoV, and SARS-CoV were given simultaneously, each at 1μg LNP-mRNA (3μg total) for both prime and boost. To generate comparable data, we started the first dose at the same day (day 0), and harvested the blood sample at the same day (day 119, i.e. 4 months, post first dose), for both sequential and mixture schedules (**Figure 5A**). We measured the antibody titers from plasma samples of both Sequential and Mixture LNP-mRNA vaccinated animals (**Figure 5B; Figure S7**). While all vaccinated animals showed certain antibody responses across all antigens tested (SARS2 WT/WA1, SARS2 Delta, SARS, MERS; both ECD and RBD), Sequential vaccination group showed significantly higher antibody responses than Mixture vaccination group across all conditions, i.e. across all antigens from these three species (**Figure 5B-C; Figure S7**). Similar with the results above, the ELISA ECD activity highly correlated with that of RBD (**Figure 5D**). We tested the neutralization activities using pseudovirus assays, which had been shown to be a robust surrogate of neutralization antibodies and widely used by the coronavirus field (e.g. see various previous literature ^31–35^). Results showed that mice in the Sequential vaccination schedule showed significantly higher neutralization activities than those in the Mixture vaccination group, and again across all three species (**Figure 5E-F**). Again, overall, such neutralization activities significantly correlated with ECD ELISA for all groups or all mice, among the spike antigens and pseudoviruses tested (**Figure S8**). These data suggested that, for LNP-mRNA vaccination against three coronavirus species under the conditions tested, vaccination in sequence can elicit more potent antibody responses than vaccination simultaneously in mixture.

**Figure 5.**
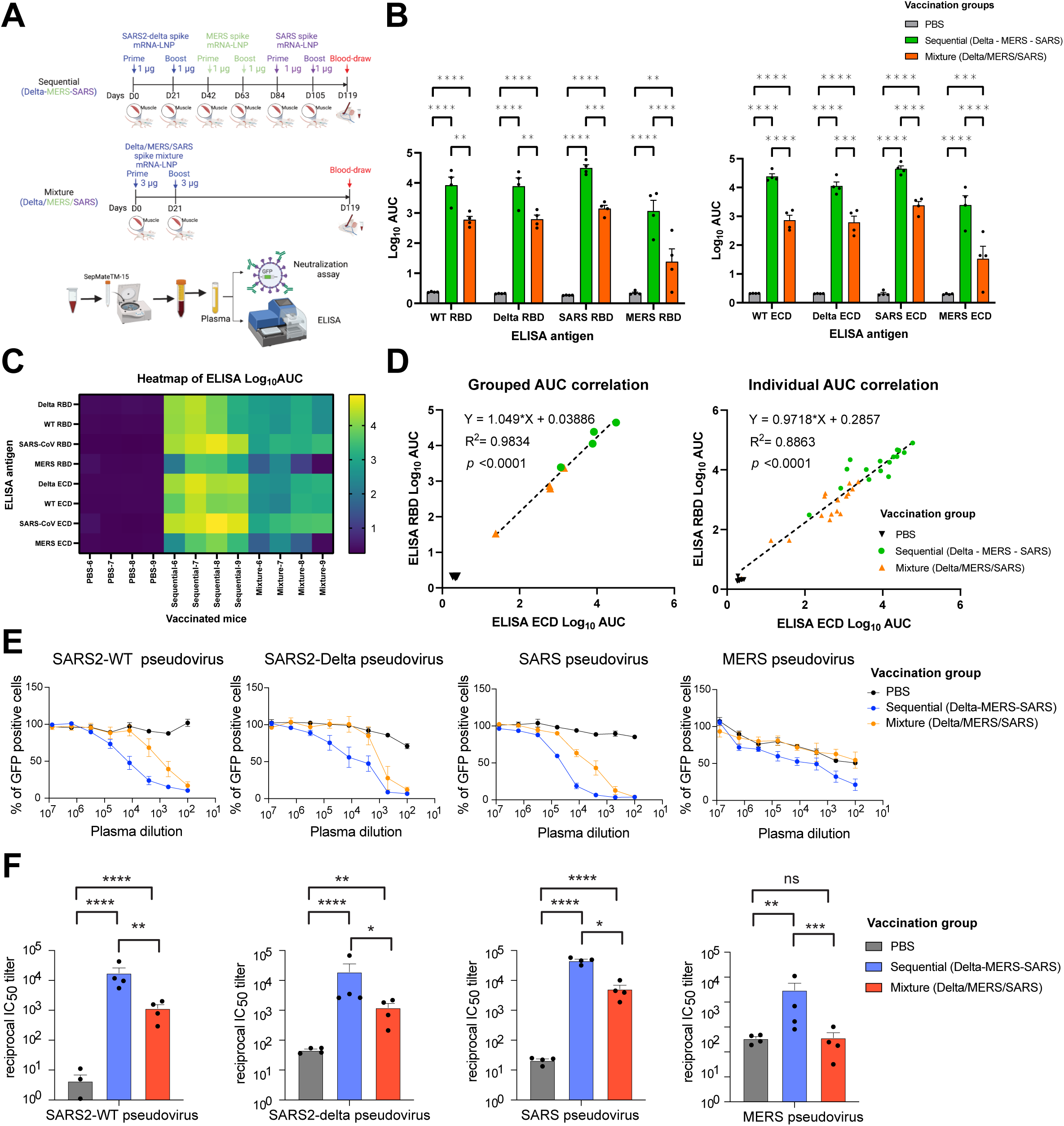
Direct comparison of sequential vs. mixture vaccination schedules against SARS-CoV-2 Delta, MERS-CoV, and SARS-CoV. **(A)** Schematics of sequential vs. mixture vaccination schedules and sampling. In the **Sequential** vaccination schedule, vaccinations of SARS-CoV-2 Delta, MERS-CoV, and SARS-CoV were given in sequence separated by 3 weeks, each with 1μg LNP-mRNA prime and 1μg LNP-mRNA boost 3 weeks apart. In the **Mixture** vaccination schedule, vaccinations of SARS-CoV-2 Delta, MERS-CoV, and SARS-CoV were given simultaneously, each at 1μg LNP-mRNA (3μg total) for both prime and boost. The first dose and the blood sample harvest were done at the same day for both sequential and mixture schedules for comparison. **(B)** Dot-box plots summarizing binding antibody titers of plasma from mice administered with PBS, Sequential or Mixture LNP-mRNA vaccinations (n = 4 each) against RBD or ECD of SARS-CoV-2 WT/WA-1 and Delta variant, as well as SARS and MERS spikes. **(C)** Heatmap of antibody titers of individual mice (one column represents one mouse) against eight spike antigens in ELISA (one row represents one antigen). **(D)** Correlation of antibody titers against RBD (y value) and ECD (x value) of same coronavirus spike, by individual mouse, or by averaged group. **(E)** Neutralization titration curves of plasma from mice treated with PBS, Sequential or Mixture LNP-mRNA vaccinations (n = 4 each); all tested against WT/WA-1 and Delta SARS-CoV-2, SARS-CoV and MERS-CoV pseudoviruses. The percent of GFP positive cells reflected the infection rate of host cells by pseudovirus and was plotted against the dilution factors of mice plasma to quantify neutralizing antibody titers. **(F)** Neutralizing antibody titers in the form of reciprocal IC50 derived from fitting the titration curves with a logistic regression model. Each dot represents data from one mouse and each group contains three mice. **Notes:** In the dot-box plots of this figure, each dot represents data from one mouse. Data are shown as mean ± s.e.m. plus individual data points in plots. Two-way ANOVA with Tukey’s multiple comparisons test was used to assess statistical significance. Statistical significance labels: * p < 0.05; ** p < 0.01; *** p < 0.001; **** p < 0.0001. Non-significant comparisons are not shown, unless otherwise noted as n.s., not significant.

## DISCUSSION

Protective vaccines are the keys to control the on-going and potential future coronavirus pandemics. Coronavirus is a group of viral species that can constantly evolve to become highly contagious and pathogenic to human. Multiple coronavirus species have emerged, and many new variants keep emerging during the spread. Pathogenic coronaviruses have emerged multiple times and infected human populations, several of which (SARS-CoV, MERS-CoV, SARS-CoV-2) have caused severe diseases and fatalities ^3, 5, 36^. Several existing less pathogenic coronavirus species (e.g. NL63, 2293, OC43, and HKU1) have been reported to have evolved hundreds to tens of thousands of years ago ^37^. Therefore, it is critical to have vaccines against multiple coronavirus species, ideally as pan-coronavirus vaccines, to help fight not only the current pandemic, but also to prevent the re-emergence of the previously existed pathogenic species, as well as constantly evolving and lurking coronavirus diseases as probable future outbreaks. Various prior efforts led to the development of SARS and MERS vaccine candidates, although at earlier stages of development ^11–15^. The COVID-19 pandemic urged an international effort for rapid development of vaccines against SARS-CoV-2 ^16^, leading to multiple successful candidates including the highly efficacious mRNA vaccines ^17, 18^. However, all these vaccines target a single species and may not offer sufficient protection against other pathogenic species. A small number of “pan-coronavirus” vaccine candidates have been recently generated and tested in animal models, using protein antigen nanocage, or mRNA encoding chimeric spike, with the focus on SARS-CoV and SARS-CoV-2 and several other non-pathogenic viruses ^38, 39^. Multiplex LNP-mRNA vaccine against the more lethal species such as SARS-CoV and MERS-CoV, also need to be rigorously tested.

Our study directly generated mRNA vaccine candidates and tested in several LNP-mRNA combinations against MERS-CoV, SARS-CoV and SARS-CoV-2, and profiled the immune responses at the single cell level.

Multiplexing of three coronavirus spike LNP-mRNA (Triplex-CoV-3µg group) produced Delta binding and neutralizing antibody titers similar to Delta LNP-mRNA alone (Delta-1µg group) at the same dose of Delta spike mRNA. Interestingly, despite of high similarity of SARS and SARS2 spike and cross reactivity of their induced antibodies, no additive effect on Delta neutralizing antibody titer was observed when combining SARS and SARS2 Delta mRNA in triplex LNP-mRNA vaccine (Triplex-CoV-3µg group vs. Delta-1µg group). When simultaneously administered with same dose of SARS, SARS2 Delta and MERS LNP-mRNA in the Triplex formulation, mice generated MERS binding and neutralizing titers lower than those of SARS and SARS2, which showed similar titer levels. The simultaneous vaccination with multiplexing may have an impact on MERS LNP-mRNA as it did for Delta LNP-mRNA as discussed above, perhaps due to immunodominance, a phenomenon well-known in immunology ^40^, although never tested in a multiplexed LNP-mRNA vaccination setting before. Our study reported the antibody responses of triplex LNP-mRNA vaccines based on MERS spike in combination with SARS and/or SARS2 Delta spikes. The level of cross-reactivity of induced antibodies was in concordance with the sequence identity between vaccine antigen and binding antigen tested in ELISA and pseudovirus assay. Two recent studies tested chimeric vaccines against multiple coronavirus species both focused Sarbecoviruses (group 2b) ^38, 39^. Different from the prior studies, the antiviral spectrum we tested here covers three highly pathogenic coronavirus species in the *Betacoronavirus* genus, and goes beyond the group 2b coronavirus category (*Sarbecoviruses*), as it includes MERS in the *Merbecovirus* subgenus. Because of low sequence similarity, the current vaccines based on SARS2 provide little protection against MERS, the most fatal coronavirus to date with a 35% mortality rate. To broaden vaccine’s anti-coronavirus spectrum, we designed and tested the triplex LNP-mRNA vaccine including SARS, SARS2 and MERS. The lower level of MERS antibody in Triplex vaccine marked a significant challenge of introducing cross-lineage antigens in multiplex vaccination.

We performed head-to-head sequential vaccination as compared to simultaneous vaccination in mixture, showing that sequential vaccination showed higher antibody responses at the endpoint. These observations and considerations may be informative for LNP-mRNA vaccination against multiple coronavirus species. In fact, this is one of the clinical precautions of vaccination, where individuals are advised to take the COVID-19 mRNA vaccine at least two weeks away from taking other vaccines. Consistent with this notion, our data with direct comparisons in animal vaccination experiments suggested that, giving the mRNA vaccine shots in sequence may be more effective in eliciting broad neutralizing antibodies than giving mRNAs simultaneously in mixture. Because of the waning immunity of coronavirus vaccines ^41, 42^, the general public, especially the immunocompromised, is recommended to receive booster shot(s) for COVID-19 vaccine. Thus, vaccination in sequence may be beneficial regardless. The direct comparison between simultaneous and sequential vaccination offers insights on optimization of vaccination schedules to provide broad and potent protective antibody immunity against three major pathogenic coronavirus species. Given that there are diverse coronavirus species with several of them being pathogenic, and many of them being potentially pathogenic in future human exposures, multiplexed vaccination against two or more species will be critical. Future design of pan-coronavirus vaccines may need seek a balance between protection breadth and depth by choosing the right number of spike antigens across coronavirus lineages. In summary, this study provided LNP-mRNA vaccine constructs designed to target SARS-CoV, SARS-CoV-2 Delta and MERS-CoV, as well as direct *in vivo* animal testing and single cell immune profiling results of multiplexed combinations as well as comparative vaccination schedules.

## Methods

### Coronavirus spike sequence alignment

The spike sequence used to produce the LNP-mRNA vaccines were aligned using Clustal Omega ^43^ and visualized in Jalview ^44^.

### Plasmid construction

The spike cDNA of SARS-CoV (Genbank accession AAP13567.1) and MERS-CoV (Genbank accession AFS88936.1) were purchased from Sino Biological (Cat # VG40150-G-N and VG40069-G-N, respectively). cDNA of SARS-CoV-2 B.1.617.2 (Delta variant) ^45^ were synthesized as gBlocks (IDT). The spike sequences were cloned by Gibson Assembly (NEB) into pcDNA3.1 plasmid for the mRNA transcription and pseudovirus assay. The plasmids for the pseudotyped virus assay including pHIVNLGagPol and pCCNanoLuc2AEGFP are gifts from Dr. Bieniasz’ lab ^46^. The C-terminal 19 (for SARS-CoV and SARS-CoV-2) or 16 (for MERS-CoV) amino acids were deleted in the spike sequence for the pseudovirus assay. To improve expression and retain prefusion conformation, six prolines (HexaPro variant, 6P) ^47^ were introduced to the SARS-CoV-2, SARS-CoV and MERS-CoV spike sequence at the homologous sites in the mRNA transcription plasmids. The furin site of SARS-CoV-2 spike (RRAR) were replaced with a GSAS short stretch to keep S1 and S2 subunits connected in the spike.

### Cell culture

HEK293T (ThermoFisher), Huh-7 and 293T-hACE2 (Dr Bieniasz’ lab) cell lines were cultured in complete growth medium, Dulbecco’s modified Eagle’s medium (DMEM; ThermoFisher) supplemented with 10% Fetal bovine serum (FBS, Hyclone), 1% penicillin-streptomycin (Gibco) (D10 media for short). Cells were typically passaged every 1-2 days at a split ratio of 1:2 or 1:4 when the confluency reached about 80%.

### In vitro mRNA transcription and vaccine formulation

Codon-optimized mRNA encoding HexaPro spikes of SARS-CoV-2 WT, Delta, SARS-CoV and MERS-CoV were synthesized *in vitro* using an Hiscribe^TM^ T7 ARCA mRNA Kit (with tailing) (NEB, Cat # E2060S), with 50% replacement of uridine by N1-methyl-pseudouridine. A linearized DNA template containing the spike open reading frame flanked by 5′ untranslated region (UTR), 3′ UTR and 3’-end polyA tail was used as for mRNA transcription. The linearization of DNA templates was achieved by digesting circular plasmids with BbsI restriction enzyme, followed by gel purification.

The mRNA was synthesized and purified by following the manufacturer’s instructions and kept frozen at -80 °C until further use. The mRNA was encapsulated in lipid nanoparticles using the NanoAssemblr^®^ Ignite™ machine (Precision Nanosystems). For the Triplex vaccine, equal mass of SARS, MERS and Delta spike mRNA were mixed before encapsulated by lipid nanoparticles. All procedures are following the guidance of manufacturers. In brief, lipid mixture was mixed with transcribed mRNA in the low pH formulation buffer 1 on Ignite instrument at a molar ratio of 6:1 (LNP: mRNA), similar to previously described ^48, 49^. The LNP encapsulated mRNA was buffer exchanged to PBS using 30kDa Amicon filter (MilliporeSigma™ UFC901024). Sucrose was added as a cryoprotectant. The particle size of mRNA-LNP was determined by DLS machine (DynaPro NanoStar, Wyatt, WDPN-06) and TEM described below. The encapsulation rate and mRNA concentration were measured by Quant-iT™ RiboGreen™ RNA Assay (ThermoFisher).

### *In vitro* mRNA expression and receptor binding validation of translated spikes

HEK293T cells were electroporated with mRNA encoding SARS, MERS or Delta spikes using Neon™ Transfection System 10 μL Kit following the standard protocol provided by manufacturer. After 12 h, the cells were collected and resuspended. To detect surface-protein expression, the cells were stained with ACE2–Fc chimera (Genscript, Z03484) or DPP4-Fc (Sino Biological, 10688-H01H) in MACS buffer (D-PBS with 2 mM EDTA and 0.5% BSA) for 30 min on ice. Thereafter, cells were washed twice and incubated with PE–anti-human FC antibody (Biolegend, 410708) in MACS buffer for 30 min on ice. Data acquisition was performed on BD FACSAria II Cell Sorter (BD). Analysis was performed using FlowJo software.

### Negative-stain TEM

5 μl of the sample was deposited on a glow-discharged formvar/carbon-coated copper grid (Electron Microscopy Sciences, catalog number FCF400-Cu-50), incubated for 1 min and blotted away. The grid was washed briefly with 2% (w/v) uranyl formate (Electron Microscopy Sciences, catalog number 22450) and stained for 1 min with the same uranyl formate buffer. Images were acquired using a JEOL JEM-1400 Plus microscope with an acceleration voltage of 80 kV and a bottom-mount 4k × 3k charge-coupled device camera (Advanced Microscopy Technologies, AMT).

### Mouse immunization

6-8 weeks old female C57BL/6Ncr (B6) mice were purchased from Charles River and used for vaccine immunogenicity study. Animals were housed in individually ventilated cages in a dedicated vivarium with clean food, water, and bedding. A maximum of 5 mice was allowed in each cage, at regular ambient room temperature (65-75°F, or 18-23°C), 40-60% humidity, and a 14 h:10 h day/night cycle. All experiments utilize randomized littermate controls. A standard two-dose schedule given 21 days apart was adopted^17^, unless otherwise noted.

For the Triplex dosage testing experiment, 1 μg Delta LNP-mRNA, 1 μg or 3 μg Triplex-CoV LNP-mRNA (equal mass mixture of Delta, MERS and SARS mRNA) were diluted to the same volume with 1X PBS and inoculated into mice intramuscularly during prime and boost.

For the Schedule comparison testing experiment, 1 μg Delta, MERS and SARS LNP-mRNA were sequentially inoculated into mice during prime and boost.

Control mice received 50µl PBS at prime and boost at the same matched time points in all experiments. In the real world setting, individuals will not receive mock mRNA-LNP formulation; and instead, in clinical trials, they receive placebo, which are usually salines (e.g. see a Moderna mRNA-1273 trial https://clinicaltrials.gov/ct2/show/NCT04796896 where Placebo is 0.9% sodium chloride (normal saline) injection. Thus, to mimic the relevant human setting and to evaluate the immune response the vaccine formulation elicits as a whole composition, PBS is chosen as a negative control to represent the placebo group or the unvaccinated.

### Sample collection, plasma and PBMCs isolation

At the defined time points, usually two weeks post the last dose of boost unless otherwise noted (e.g. day 35, or day 119, as noted in the schematics), blood was retro-orbitally collected from mice. The PBMCs and plasma were isolated from blood via SepMate-15 (StemCell Technologies). 200 µl blood was immediately diluted with 800 ul PBS with 2% FBS. The diluted blood was then added to SepMate-15 tubes with 5ml Lymphoprep (StemCell Technologies). 1200 x g centrifugation for 20 minutes was applied to isolate RBCs, PBMCs and plasma. 200ul diluted plasma was collected from the surface layer. Then the solution at the top layer containing PBMCs was poured to a new tube. PBMCs were washed once with PBS + 2% FBS before being used in downstream analysis. The separated plasma was used in ELISA and neutralization assay. PBMCs were collected for single cell profiling using a 10xGenomics platform.

### ELISA

The 384-well ELISA plates were coated with 3 μg/ml of antigens overnight at 4 degree. The antigen panel used in the ELISA assay includes SARS-CoV-2 spike S1+S2 ECD and RBD of 2019-nCoV WT (Sino Biological, ECD 40589-V08B1 and RBD 40592-V08B), Delta variant B.1.617.2 (SINO, ECD 40589-V08B16 and RBD 40592-V08H90), SARS-CoV (ECD Sino Biological 40634-V08B and RBD Fisher 50-196-4017) and MERS-CoV (ECD Sino Biological 40069-V08B and RBD Fisher 50-201-9463). Plates were washed with PBS plus 0.5% Tween 20 (PBST) three times using the 50TS microplate washer (Fisher Scientific, NC0611021) and blocked with 0.5% BSA in PBST at room temperature for one hour. Plasma was serially diluted twofold or fourfold starting at a 1:500 dilution. Samples were added to the coated plates and incubate at room temperature for one hour, followed by washes with PBST five times. Anti-mouse secondary antibody (Fisher, Cat# A-10677) was diluted to 1:2500 in blocking buffer and incubated at room temperature for one hour. Plates were washed five times and developed with tetramethylbenzidine substrate (Biolegend, 421101). The reaction was stopped with 1 M phosphoric acid, and OD at 450 nm was determined by multimode microplate reader (PerkinElmer EnVision 2105). The binding response (OD450) were plotted against the dilution factor in log_10_ scale to display the dilution-dependent response. The area under curve of the dilution-dependent response (Log_10_ AUC) was calculated to evaluate the potency of the serum antibody binding to spike antigens.

### Pseudovirus neutralization assay

HIV-1 based SARS-CoV-2 WT, B.1.617.2 (delta) variant, SARS and MERS pseudotyped virions were generated using corresponding spike sequences, and applied in neutralization assays. The pseudotyped virus was packaged using a coronavirus spike plasmid, a reporter vector and a HIV-1 structural protein expression plasmid. The reporter vector, pCCNanoLuc2AEGFP, and HIV-1 structural/regulatory proteins (pHIVNLGagPol) expression plasmid were from Bieniasz lab. The spike plasmid for SARS-CoV-2 WT pseudovirus truncated 19 C-terminal amino acids of S protein (SARS-CoV-2-Δ19) and was from Bieniasz lab. Spike plasmids expressing C-terminally truncated SARS-CoV-2 B.1.617.2 variant S protein (Delta variant-Δ19), SARS-CoV S protein (SARS-CoV-Δ19) and MERS S protein (MERS-CoV-Δ16) were generated based on the pSARS-CoV-2-Δ19. Briefly, 293T cells were seeded in 150 mm plates, and transfected with 21 µg pHIVNLGagPol, 21 µg pCCNanoLuc2AEGFP, and 7.5 µg of corresponding spike plasmids, in the presence of 198 µl PEI. At 48 h after transfection, the 20-ml supernatant was harvested and filtered through a 0.45- μm filter, and concentrated before aliquoted and frozen in -80°C.

The SARS-CoV and SARS-CoV-2 pseudovirus neutralization assays were performed on 293T-hACE2 cell, while the MERS-CoV neutralization assay was performed on Huh-7 cells. One day before infection, 293T-hACE2 cells were plated in a 96 well plate with 0.01 x10^6^ cells per well. In the next day, plasma collected from PBS or LNP-mRNA immunized mice were 5-fold serially diluted with complete growth medium starting from 1:100. 55 μL aliquots of diluted plasma were mixed with the same volume of SARS-CoV-2 WT, Delta variant, SARS or MERS pseudovirus. The mixture was incubated for 1 hr in the 37 °C incubator, supplied with 5% CO2. Then 100 μL of mixtures were added into 96-well plates with 293T-hACE2 or Huh-7 cells. Plates were incubated at 37°C for 48 hr. Then host cells were collected and the percent of GFP-positive cells were analyzed with Attune NxT Acoustic Focusing Cytometer (ThermoFisher). The 50% inhibitory concentration (IC50) was calculated with a four-parameter logistic regression using GraphPad Prism (GraphPad Software Inc.). If the curve of individual mouse fails to produce positive fit (i.e. negative titer), suggestive of no neutralization activity, the value was converted to zero.

### Correlation analysis

Correlation analysis of ELISA, pseudovirus neutralization and authentic virus neutralization data were performed using the respective data collected. Linear regression model was used to evaluate the correlations between ELISA RBD and ECD AUCs, pseudovirus neutralization and authentic virus neutralization log10 IC50. Model fitting and statistical analysis were performed in Graphpad Prism9.1.2. Correlations of data points from either individual mouse, or group average of different vaccination groups, were analyzed separately. The vaccination-group ELISA AUC or neutralization log10 IC50 were calculated from the average of individual value in each group. Due to assay-dependent PBS background level, only non-PBS data points were included in the correlation analysis.

### Single cell RNA-seq

PBMCs were collected from mRNA-LNP vaccinated and control mice were collected as described above for mouse immunization and sample collection, and normalized to 1000 cells/μL. Standard volumes of cell suspension were loaded to achieve targeted cell recovery to 10000 cells. The samples were subjected to 14 cycles of cDNA amplification. Following this, gene expression (GEX) libraries were prepared according to the manufacturer’s protocol (10x Genomics). All libraries were sequenced using a NovaSeq 6000 (Illumina) with 2*150 read length.

### Single cell data analysis for immune repertoire profiling and transcriptomic signatures

Both standard established pipelines and custom scripts were used for processing and analyzing single cell GEX data. Illumina sequencing data was processed using the Cellranger v6.0.1 (10x Genomics) pipeline, aligning reads to the mm10 reference transcriptome and aggregating all samples. Cellranger outputs were then preprocessed using a modified Seurat v4.0.5 workflow with the R statistical programming language^50^. Briefly, individual sample data sets were filtered for quality cells (200-2000 RNA features and < 5% mitochondrial RNA), log-normalized, scaled, and quality features were selected to calculate low-dimensional “anchors” (reciprocal-PCR dimensional reduction, k = 20, anchors = 2000), which were used to integrate the different sample data sets^51^. Integrated single-cell data were scaled, centered, clustered by shared nearest neighbors graph (k = 20, first 12 PCA dimensions, chosen by the elbow plot method) with modularity optimization (Louvain algorithm with multilevel refinement, empirically chosen resolution = 0.31). Clustered cells were visualized in low-dimensional space by uniform manifold approximation and projection (UMAP; first 12 PCA dimensions) ^52^, and clusters were labeled as immune cell types via canonical marker expression, based on scaled-mean expression and expression detection rate for the cluster. Immune cell subtypes were identified for B cells, plasma cells, activated CD4 T cells, and mononuclear myeloid cells by sub-setting the cells of each group, rescaling with mt-RNA % as a covariate, centering, UMAP dimensional reduction as before (first 14, 11, 16, and 10 PCA dimensions for B cells, plasma cells, activated CD4 T cells, and myeloid cells, respectively), and clustering was performed as previously described (empirically chosen modularity resolution = 0.20, 0.10, 0.25, and 0.10 for B cells, plasma cells, activated CD4 T cells, and myeloid cells, respectively), but canonical marker genes were used as features. To show that the cell type populations displayed distinct transcriptional profiles, markers were identified for each cluster vs all other cells using Wilcoxon rank sum testing of scaled data (SeuratWrappers::RunPrestoAll R function), while down-sampling to 5000 cells per cluster. The top 10 mean log fold change genes were selected from each cell type to visualize by heatmap with hierarchical clustering.

Differential expression was performed using the edgeR analysis pipeline and quasi-likelihood (QL) F tests ^53, 54^. Specifically, raw single-cell expression data was filtered to include genes with > 5% detection rate across all cells, genes were TMM-normalized, fitted to a QL negative binomial generalized linear model using trended dispersion estimates with cell detection rate and treatment as covariates, and empirical Bayes QL F tests were performed with treatment as the coefficient equal to zero under the null hypothesis ^54^.

Pathway enrichment analyses were performed for differentially expressed genes (DEG; absolute log2(x+1) expression fold-change > 0.5, FDR-adjusted p value (q) < 0.01) using the gost function of the gProfiler2 R package ^55, 56^ with biological process gene ontologies (GO) for mus musculus, an adjusted p value-ordered gene list, and known genes as the domain for the statistics. In addition, the analysis p values were adjusted for multiple testing using the gProfiler gSCS method. Results were filtered to include GO terms <=600 genes in size that intersected > 2 DEG, an absolute activation score (mean log2(x+1) expression fold change of GO term DEGs) > 0.5, and an adjusted p < 0.01. Network analyses were performed by (1) creating network graphs with filtered pathway results as nodes and GO term similarity coefficients as edges (coefficients = 50% jaccard + 50% overlap scores; edge similarity threshold = 0.375), (2) finding graph clusters via the Leiden algorithm using the modularity method with similarity coefficients as weights (resolution = 0.5, iterations = 1000), and (3) labeling clusters by their most significant GO term (meta-pathway). Meta-pathway genes were visualized by heatmap, using log-normalized, scaled expression for GO term genes that were differentially expressed in vaccination groups compared to the PBS control. Custom R scripts were used for generating various plots.

### Standard statistics

Standard statistical methods were applied to non-high-throughput experimental data. The statistical methods are described in figure legends and/or supplementary Excel tables. The statistical significance was labeled as follows: n.s., not significant; * p < 0.05; ** p < 0.01; *** p < 0.001; **** p < 0.0001. Prism (GraphPad Software) and RStudio were used for these analyses. Additional information can be found in the supplemental excel tables.

### Schematic illustrations

Schematic illustrations were created with Affinity Designer or BioRender.

### Replication, randomization, blinding and reagent validations

Replicate experiments have been performed for all key data shown in this study.

Biological or technical replicate samples were randomized where appropriate. In animal experiments, mice were randomized by littermates.

Regular experiments were not blinded. NGS data processing were blinded using metadata. Subsequent analyses were not blinded.

Commercial antibodies were validated by the vendors, and re-validated in house as appropriate. Custom antibodies were validated by specific antibody - antigen interaction assays, such as ELISA. Isotype controls were used for antibody validations.

Cell lines were authenticated by original vendors, and re-validated in lab as appropriate. All cell lines tested negative for mycoplasma.

## Supplementary Materials

Figs. S1 to S8 Data files S1 to S2

## Supplemental Datasets

### Dataset 1 | Single cell GEX of multiplexed LNP-mRNA vaccinated animals

Tabs in this dataset:

- Metadata of merged single cell GEX dataset.
- Clustering of scGEX dataset.
- Two-dimensional UMAP embeddings.
- Wilcoxon statistics for cluster-specific differentially expressed genes
- Markers for clustering immune cell subsets.
- Statistics for cell type proportions across treatment groups.
- Results of the differential expression (DE) analyses of treatment vs PBS groups in different cell types.
- Results of the differential expression (DE) analyses of Triplex vs Delta treatment groups in different cell types.
- Results of the gProfiler pathway analysis in different comparisons.

### Dataset 2 | Supplemental Source Data and Statistics of non-NGS experiments

Supplemental excel file(s) contains all original data and statistics for non-NGS experiments.

## Acknowledgments

We thank various members from Chen and Wilen labs for discussions and support. We thank staffs from various Yale core facilities (Keck, YCGA, HPC, YARC, CBDS and others) for technical support. We thank Drs. Tsemperouli, Karatekin, Lin, Wang, Castaldi and others for providing equipment and related support. We thank various support from Department of Genetics; Institutes of Systems Biology and Cancer Biology; Dean’s Office of Yale School of Medicine and the Office of Vice Provost for Research.

## Institutional Approval

This study has received institutional regulatory approval. All recombinant DNA (rDNA) and biosafety work were performed under the guidelines of Yale Environment, Health and Safety (EHS) Committee with approved protocols (Chen-15-45, 18-45, 20-18, 20-26). All animal work was performed under the guidelines of Yale University Institutional Animal Care and Use Committee (IACUC) with approved protocols (Chen-2018-20068; Chen-2020-20358; Chen 2021-20068; Wilen 2021-20198).

## Author contributions

Conceptualization: SC

Methodology: LP, ZF, PAR, AM, JJP, QL, XZ, MBD, CBW, SC

Investigation: LP, ZF, PAR, AM, JJP, QL, XZ, MBD, CBW, SC

Visualization: LP, ZF, PAR, AM, JJP, BZ

Funding acquisition: SC, CBW

Project administration: SC, CBW

Supervision: SC, CBW, HZ

Writing – original draft: LP, ZF, PAR, SC

Writing – review & editing: LP, ZF, PAR, CBW, SC

## Data and materials availability

All data generated or analyzed during this study are included in this article and its supplementary information files. Specifically, source data and statistics for non-high-throughput experiments are provided in a supplementary table excel file. High-throughput experiment data are provided as processed quantifications in Supplemental Datasets. Genomic sequencing raw data are deposited to Gene Expression Omnibus (GEO) with a pending accession code. Codes that support the findings of this research are being deposited to a public repository such as GitHub. Additional information related to this study are available from the corresponding authors upon reasonable request.

## Supplemental Figure legends

**Supplemental Figure 1.**
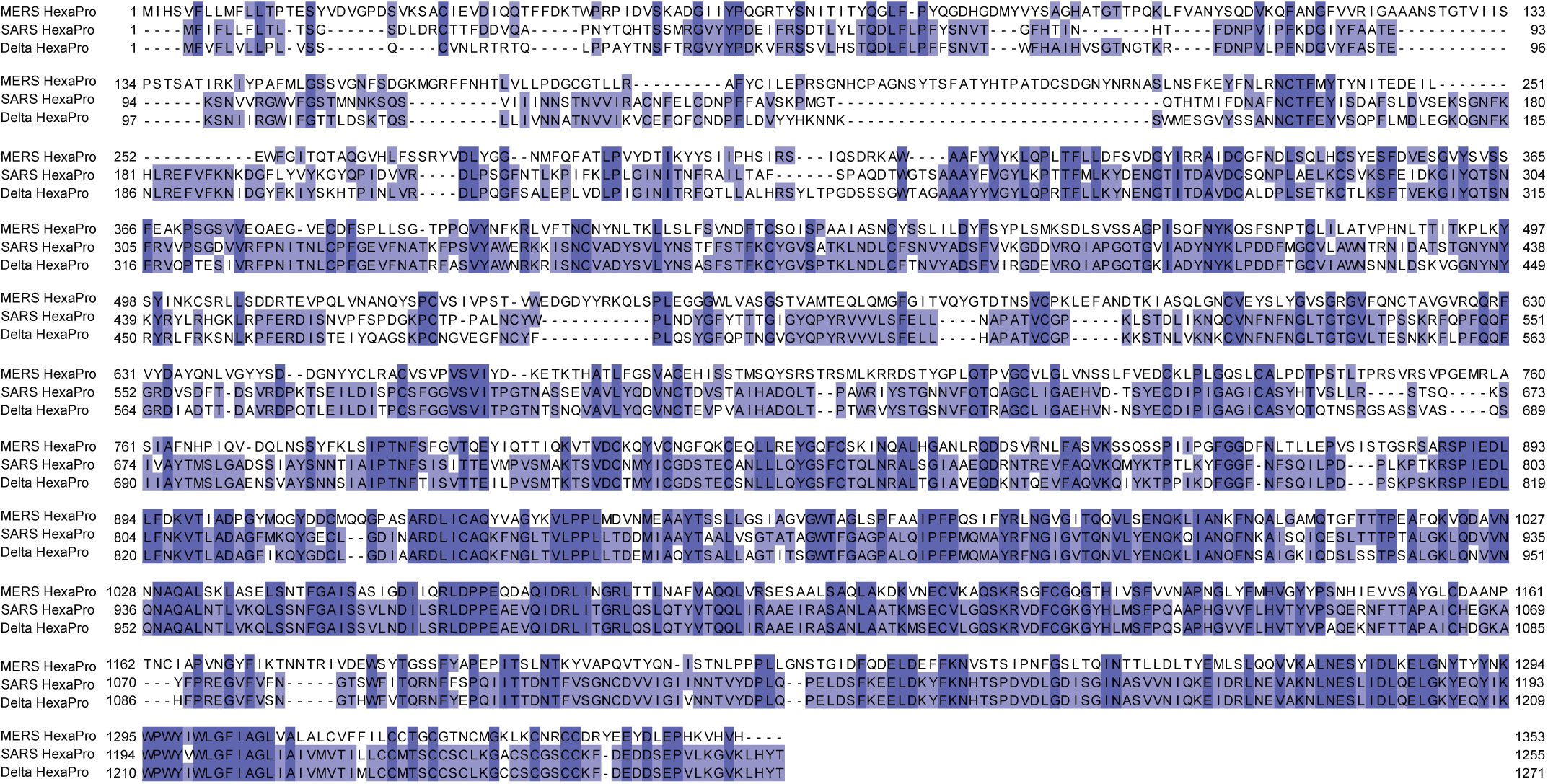
(Figure S1) Sequence analysis of engineered mRNA-encoded spike proteins of three pathogenic human coronavirus species. Sequence alignment of spikes of SARS-CoV-2 Delta variant, SARS-CoV and MERS-CoV used in the LNP-mRNA vaccine. The full-length spike sequences of these three pathogenic human coronavirus species were aligned and their degree of identity at each residue was color coded by a gradient blue color.

**Supplemental Figure 2.**
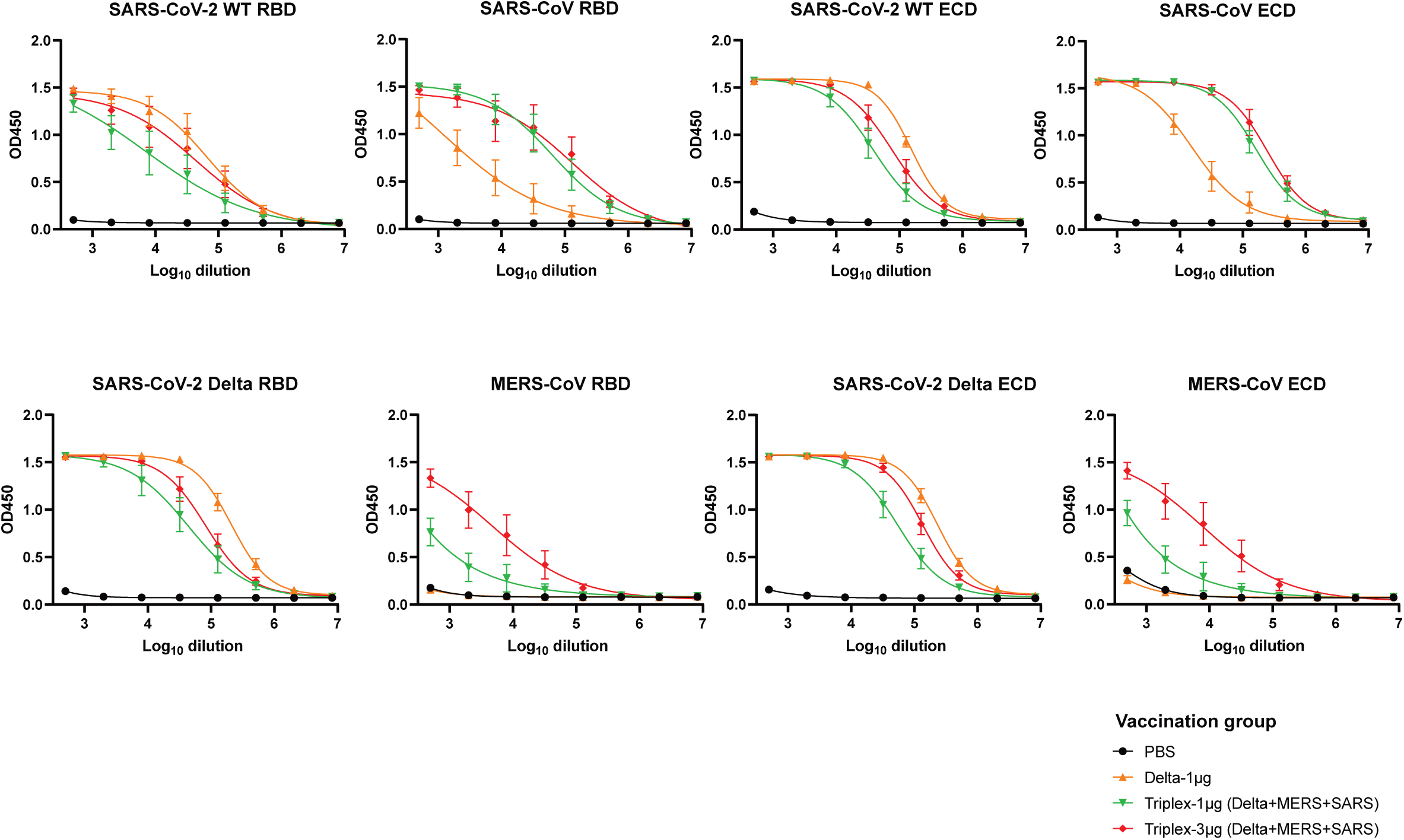
(Figure S2) ELISA titration of RBD and ECD antigens. Two types (RBD top panel and ECD bottom panel) of ELISA spike antigens derived from SARS-CoV-2 WT, Delta, SARS and MERS were used to evaluate the potency of binding antibodies induced by LNP-mRNA vaccines. The mice were intramuscularly injected with two doses (x2, 2 weeks apart) of PBS, 1μg SARS-CoV-2 Delta variant LNP-mRNA (delta), 1μg or 3μg equal mass mixture of Delta, SARS and MERS mRNA delivered by LNP (Triplex-CoV).

**Supplemental Figure 3.**
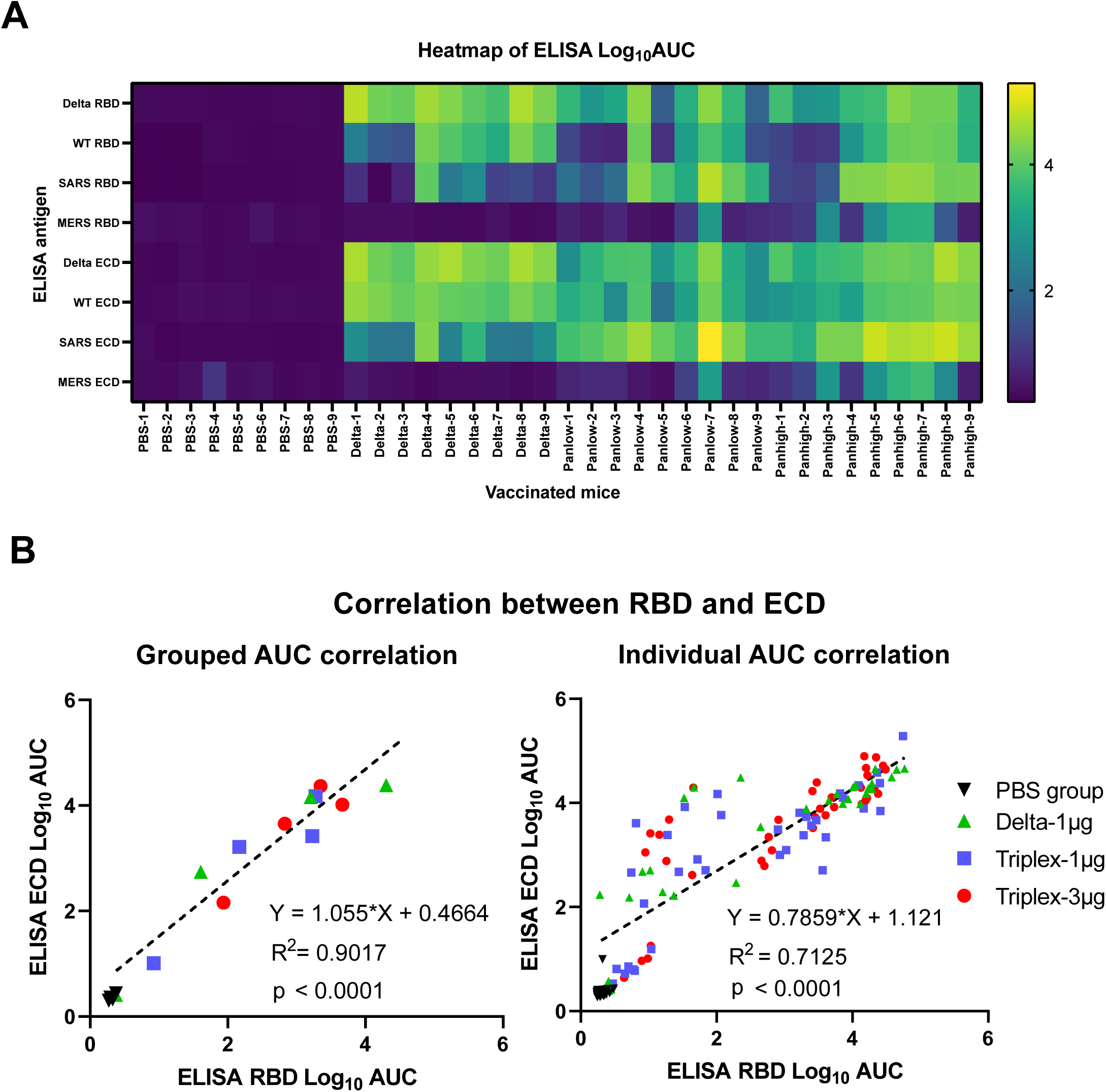
(Figure S3) Overall ELISA heatmap and correlation analysis of plasma samples from mice treated with PBS or different LNP-mRNA vaccines. **(A)** Overall heatmap of antibody titers of individual mice (one column represents one mouse) against eight spike antigens in ELISA (one row represents one antigen). **(B)** Correlation of antibody titers against RBD (y value) and ECD (x value) of same coronavirus spike, by individual mouse, or by averaged group.

**Supplemental Figure 4.**
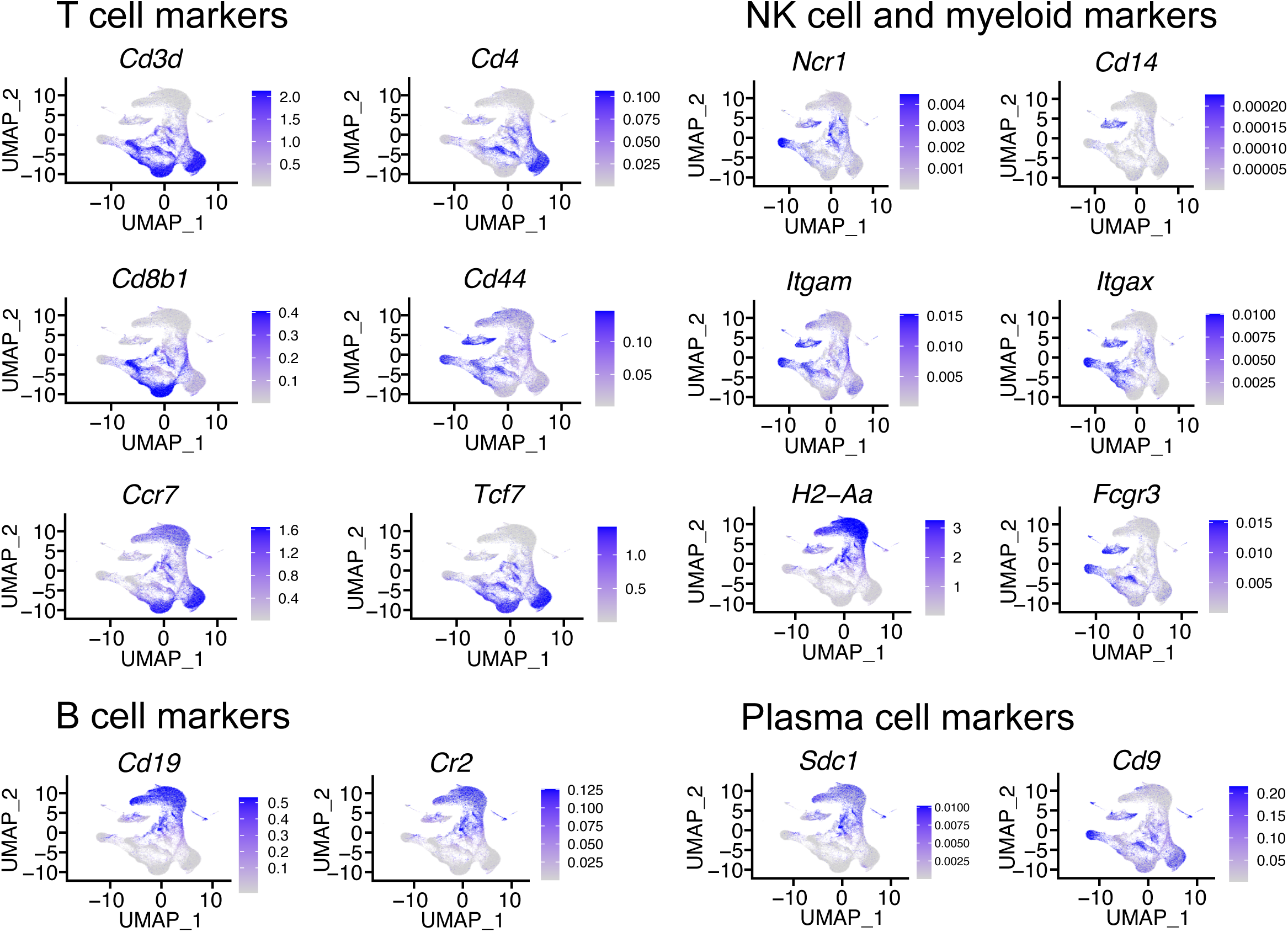
(Figure S4) Single cell transcriptomics visualization, clustering and cell type identification. UMAP visualization, colored by the scaled expression of representative cell type-specific markers in T cells, NK cells, myeloid cells, B cells, and plasma cells.

**Supplemental Figure 5.**
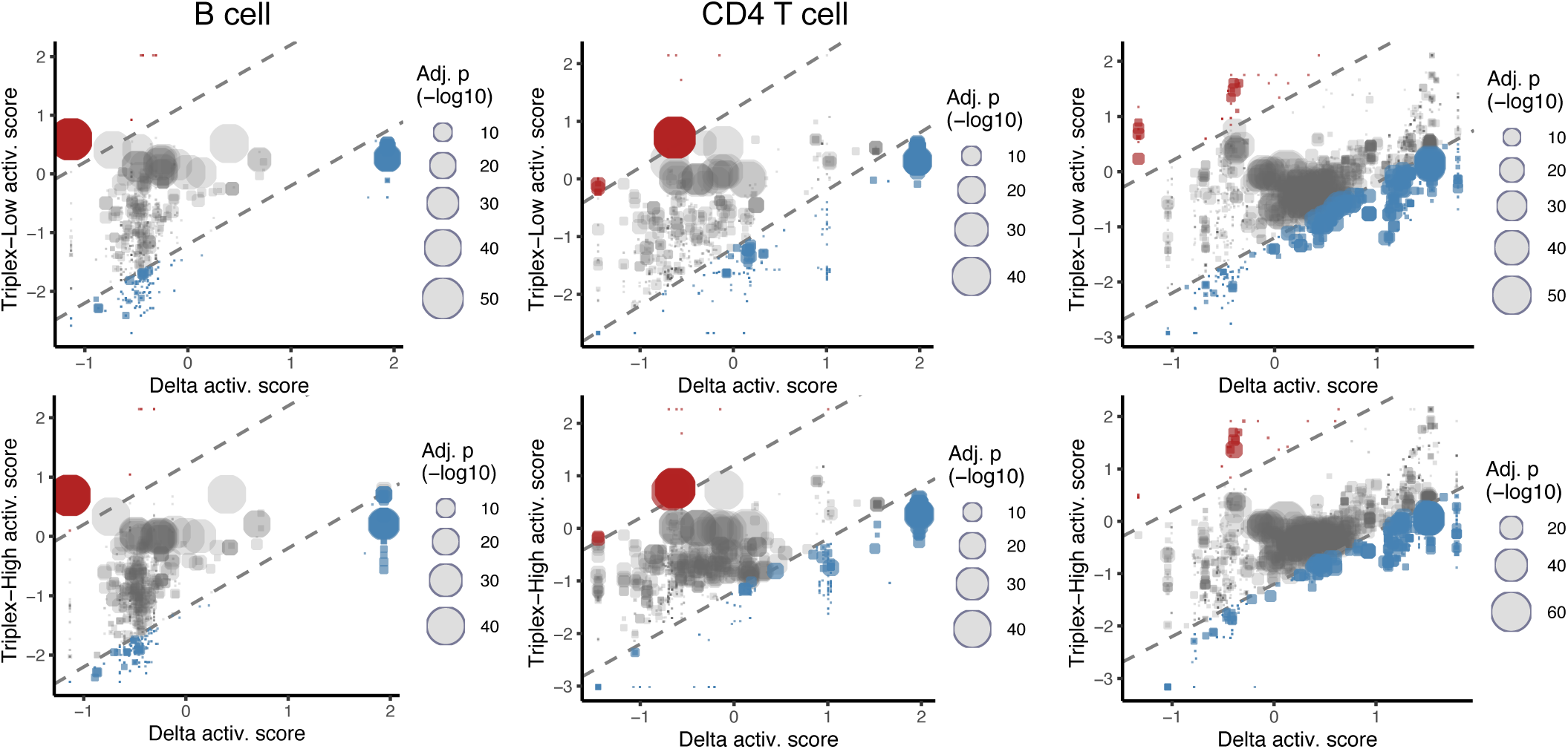
(Figure S5) Additional pathway analysis of differentially expressed genes compared between vaccination groups in different cell types in the single cell RNA-seq data. Bubble plots of overall biological process pathways of differentially expressed genes compared between vaccination groups in different cell types.

**Supplemental Figure 6.**
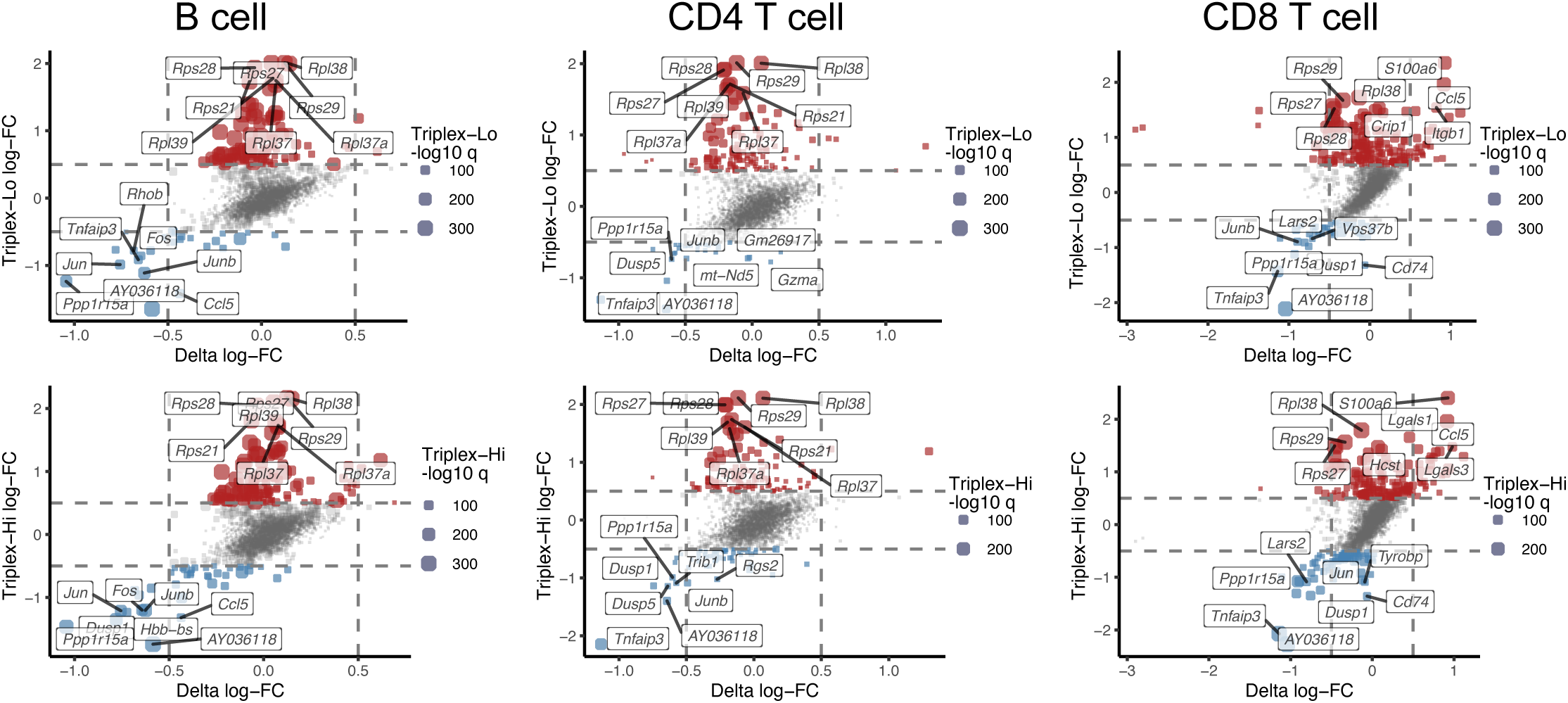
(Figure S6) Differential expression, pathway signature and gene set cluster analyses of single cell transcriptomics for animals vaccinated by multiplexed LNP-mRNAs. Square plots compare differential expression (DE) of mixCoV-vs-PBS (y axis) to Delta-vs-PBS (x axis) analyses. Each gene is presented by a dot, positioned by the log2(x+1) fold change in either DE analysis and sized by the -log10 FDR-adjusted p value. Genes that are upregulated or down regulated in mixCoV-vs-PBS are shown as red or blue dots, respectively. Analyses were done for B cell, CD4 T cell and CD8 T cell populations.

**Supplemental Figure 7.**
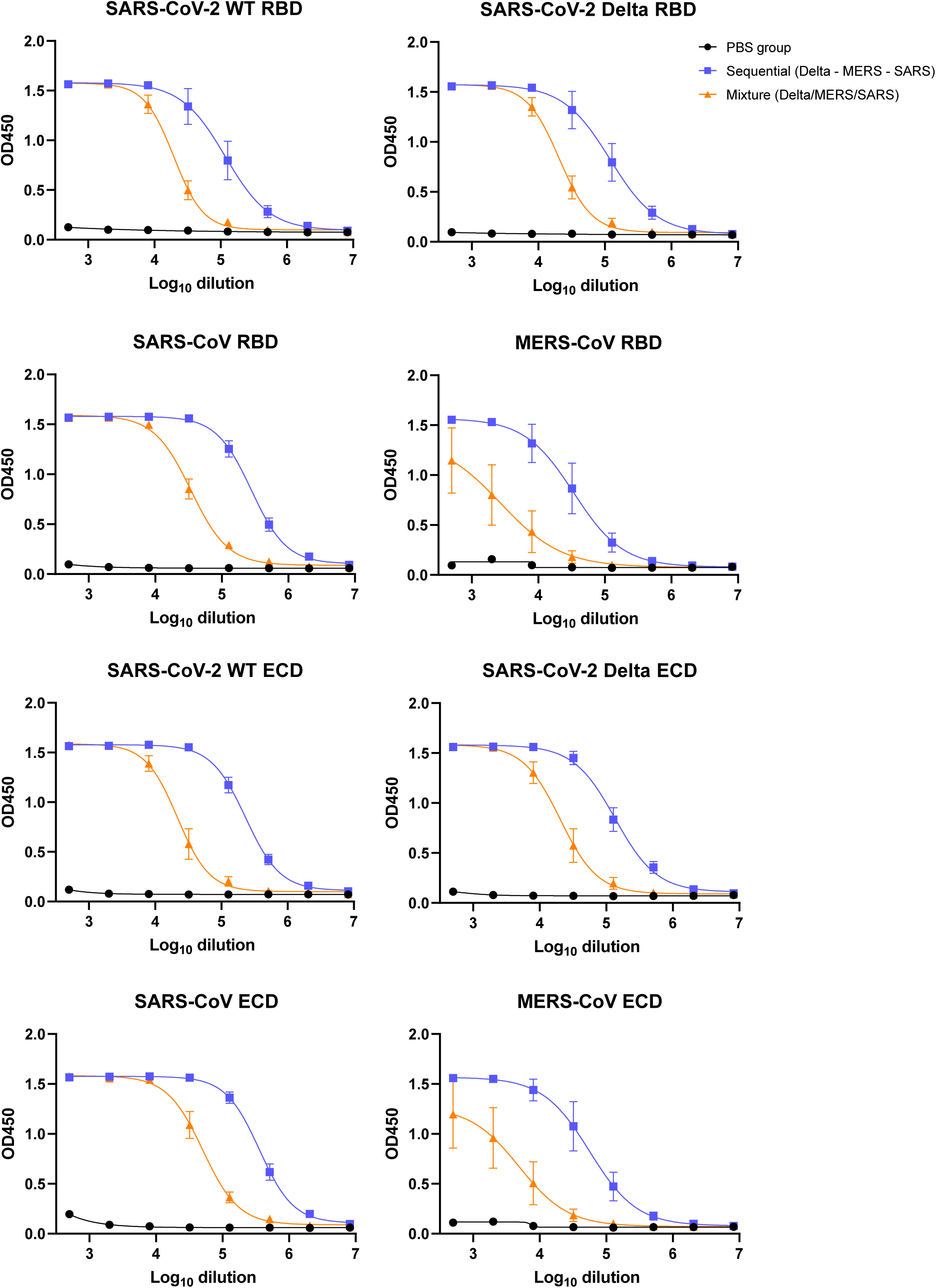
(Figure S7) ELISA OD450 titration curves over serial log10-transformed dilution points of plasma from mice treated with PBS, sequential or mixture LNP-mRNA vaccinations. The Sequential vaccination mice were intramuscularly injected with two doses (x2, 3 weeks between prime and boost) of 1μg SARS-CoV-2 Delta, MERS, SARS LNP-mRNA, three weeks apart, in this sequence (Sequential Delta-MERS-SARS). The Mixture vaccination mice were intramuscularly injected with two doses (3 weeks between prime and boost) 3μg equal mass mixture (1μg each) of Delta, SARS and MERS LNP-mRNA (Mixture Delta/MERS/SARS).

**Supplemental Figure 8.**
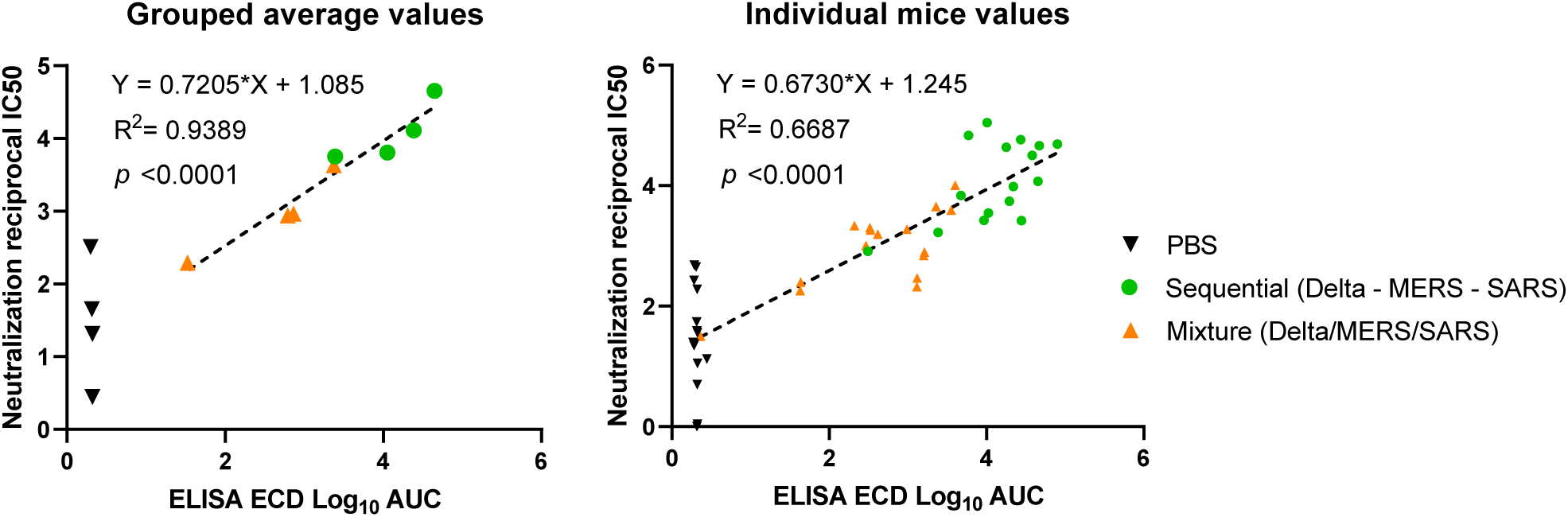
(Figure S8) Direct comparison of sequential vs. mixture vaccination schedules against SARS-CoV-2 Delta, MERS-CoV, and SARS-CoV. **(A)** Heatmap of antibody titers of individual mice (one column represents one mouse) against eight spike antigens in ELISA (one row represents one antigen). **(B)** Correlation of antibody titers against RBD (y value) and ECD (x value) of same coronavirus spike, by individual mouse, or by averaged group. **(C)** Neutralization titration curves of plasma from mice treated with PBS, Sequential or Mixture LNP-mRNA vaccinations (n = 4 each); all tested against WT/WA-1 and Delta SARS-CoV-2, SARS-CoV and MERS-CoV pseudoviruses. The percent of GFP positive cells reflected the infection rate of host cells by pseudovirus and was plotted against the dilution factors of mice plasma to quantify neutralizing antibody titers. **(D)** Correlation of neutralization IC50 vs. antibody titers against ECD of same coronavirus spike, by individual mouse, or by averaged group.

